# Catecholamine Challenge Uncovers Distinct Mechanisms for Direct Versus Indirect, but Not Social Versus Non-Social, Learning

**DOI:** 10.1101/303982

**Authors:** Jennifer L. Cook, Jennifer C. Swart, Monja I. Froböse, Andreea Diaconescu, Dirk E. M. Geurts, Hanneke E. M. den Ouden, Roshan Cools

**Affiliations:** School of Psychology, University of Birmingham, Edgbaston, Birmingham, United Kingdom; Donders Institute for Brain, Cognition and Behaviour, Radboud University, Nijmegen, The Netherlands; Translational Neuromodeling Unit, Institute for Biomedical Engineering, University of Zurich & ETH Zurich, Switzerland; Department of Psychiatry, Radboud University Medical Centre, Nijmegen, The Netherlands

## Abstract

Evidence that social and individual learning are at least partially dissociable sustains the belief that humans possess adaptive specialisations for social learning. However, in most extant paradigms, social information comprises an indirect source that can be used to supplement one’s own, direct, experience. Thus, social and individual learning differ both in terms of social nature (social versus non-social) and directness (indirect versus direct). To test whether the dissociation between social and individual learning is best explained in terms of social nature or directness, we used a catecholaminergic challenge known to modulate learning. Two groups completed a decision-making task which required direct learning, from own experience, and indirect learning from an additional source. The groups differed in terms of whether the indirect source was social or non-social. The catecholamine transporter blocker, methylphenidate, affected direct learning by improving adaptation to changes in the volatility of the environment but there was no effect of methylphenidate on learning from the social or non-social indirect source. Thus, we report positive evidence for a dissociable effect of methylphenidate on direct and indirect learning, but no evidence for a distinction between social and non-social. These data fail to support the adaptive specialisation view, instead providing evidence for distinct mechanisms for direct versus indirect learning.

---

The last decade has seen a burgeoning interest in studying the neural and computational mechanisms that underpin social learning, our ability to learn from other people. Notable findings in this field include the observation that the same computations, based on the calculation of prediction error, are involved in social learning and learning from personal experiences of reward (individual learning)^1^; both social and individual learning have been associated with activity in common brain regions^2^; and, prior preferences bias social learning as they do individual learning^3^. Such findings promote the view that ‘domain-general’ learning mechanisms underpin social learning^4,5^: we learn from other people in the same way that we learn from any other stimulus in our environment.

Though this conclusion may be of little surprise to the neuroscientist, it is likely to raise the eyebrows of many evolutionary psychologists, behavioural ecologists and others studying the evolution of social learning. In attempting to explain how rich, cumulative, human culture has come about, academics in these fields have argued that humans possess social-specific learning mechanisms – adaptive specialisations moulded by natural selection to cope with the pressures of group living^6,7^. According to this view, these adaptive specialisations differ from the domain-general associative processes that animals use to learn about their environment. Thus, evidence that social learning depends on domain-general mechanisms, is at odds with the adaptive specialisation view.

Despite evidence to the contrary, the view that there is something special about *social* learning persists. This view is implicit in neuroscience’s continued examination of the neural and computational mechanisms of social learning: if it were a foregone conclusion that all instances of social learning can be explained by domain-general principles there would be no mileage in further investigations of specifically social learning. Consistent with the hypothesis that social learning is special, studies have demonstrated that social and individual learning are at least partially dissociable. Behrens and colleagues^8^ demonstrated activity relating to social and individual learning in dissociable neural pathways, and a number of studies have found correlations between personality traits and social, but not individual, learning^9–11^.

Studies, such as those mentioned above, that have dissociated social and individual learning have typically used a single task wherein both social and individual learning happen concurrently. Though this presents advantages in terms of ecological validity (everyday decisions often require the integration of social information with information derived from one’s own experience), it also presents a confound: in this situation the social information (e.g. social advice) comprises an indirect, additional, source of information that should be integrated with one’s own, direct, experience. For example, in the Behrens et al. study participants were required to choose between a blue and a green box to accumulate points. Outcome information came in the form of a blue or green indicator thus *directly* informing participants about whether they had made the correct choice on the current trial (i.e. if the outcome indicator was blue, then the blue box was correct). Each trial also featured a red frame, which represented social information, surrounding one of the two boxes. The outcome indicator *indirectly* informed participants about the veracity of the frame: if the outcome was blue AND the frame surrounded the blue box, then the frame was correct. Thus, in most extant paradigms, social and individual learning differ both in terms of social nature (social or non-social) and directness (indirect or direct). Consequently, it is unclear which of these factors accounts for the previously observed dissociations between social and individual learning. Notably, a social-nature-based explanation would support the adaptive specialisation view, whereas a directness-based explanation would not.

Here we used a psychopharmacological challenge to test whether the dissociation between social and individual learning is best explained in terms of the social versus non-social or the indirect versus direct nature of the learning source. We employed a between-groups design wherein both groups completed a modified version of the decision-making task employed by Behrens and colleagues in which participants could improve their performance by learning from their own previous experience, and from an additional indirect source of information. For one group the indirect source was social in nature (Social Group). For the Non-Social Group the indirect source comprised a system of rigged roulette wheels (explained below) and was thus non-social in nature. All other aspects of the task including learning schedules and visual inputs were identical across the groups. Participants completed the task once under placebo (PLA) and once after 20mg of the catecholamine transporter blocker methylphenidate (MPH; order counterbalanced).

Data were analysed a) in terms of the commonly used learning indices win-stay and lose-shift scores, and b) by fitting a mathematical model of learning to estimate learning rates. Electrophysiological studies of non-human animals, human pharmacological imaging studies, and theoretical and modelling work have linked the catecholamines dopamine^12–14^ and noradrenaline^15,16^ to various learning indices including win-stay, lose-shift scores and learning rates. By blocking the reuptake of catecholamines, MPH prolongs the effects of catecholamine release^17^. Thus we predicted that these measures would be sensitive to our drug manipulation, consistent also with a recent randomised control trial with healthy adults demonstrating that MPH increased learning rates during performance of a probabilistic learning task^18^. Our crucial question, however, was whether the effect of MPH on learning varied as a function of the social versus non-social or, rather, the direct versus indirect nature of the learning source. A pattern of data wherein the Social and Non-social Groups exhibit comparable differences in the effect of MPH on direct and indirect learning would fail to support the adaptive specialisation view and would instead suggest that dissociable effects of MPH are driven by indirect versus direct learning. Alternatively, if the effect of MPH were to differ between Social and Non-social Groups, then this would strengthen the adaptive specialisation view.

## RESULTS

102 participants took part on two separate test days. The study followed a double-blind cross-over design such that 50% of participants received 20mg MPH on day 1 and PLA on day 2, for 50% of participants the order was reversed. On each test day, participants completed a complex probabilistic learning task (**Figure 1**)^8,10^. On each trial participants chose between a blue and a green stimulus to accumulate points. The probability of reward associated with the blue and green stimuli varied according to a probabilistic schedule (see **Supplementary Information 1**). Mimicking the complexity of real life decision making, all probabilistic schedules comprised stable phases in which the probability of reinforcement was held constant for between 30 and 60 trials, and volatile phases in which probabilities changed every 10 – 20 trials. Participants could use their personal experience of the reward history to track these changing probabilities (direct learning). Importantly, on each trial, before participants made a response they saw a frame surrounding one of the two options. Participants were randomly allocated to two experimental groups, ‘Social’ or ‘ Non-social’. Participants in the Social Group were informed that the frame represented the most popular choice selected by a group of participants who previously played the task. Importantly, the utility of this information followed one of four different probabilistic schedules that participants could track throughout the course of the experiment. As with the blue and green boxes, the probabilistic schedule underpinning the utility of the frame comprised stable phases in which the probability that the frame surrounded the correct option was held constant, and volatile phases in which probabilities changed every 10 – 20 trials. Participants were informed that the social information had been ‘mixed up’ such that … “you could be in a phase where the group’s information is pretty useful … but be careful, this could change! Sometimes you will see less useful information – for example from the beginning of their experiment where they didn’t have a very good idea of what was going on”. Participants in the Non-social Group were instructed that the red frame represented the outcome from a system of rigged virtual roulette wheels which fluctuated between providing predominantly correct and predominantly uninformative suggestions (see **Supplementary Information 2** for instruction scripts).

**Figure 1.**
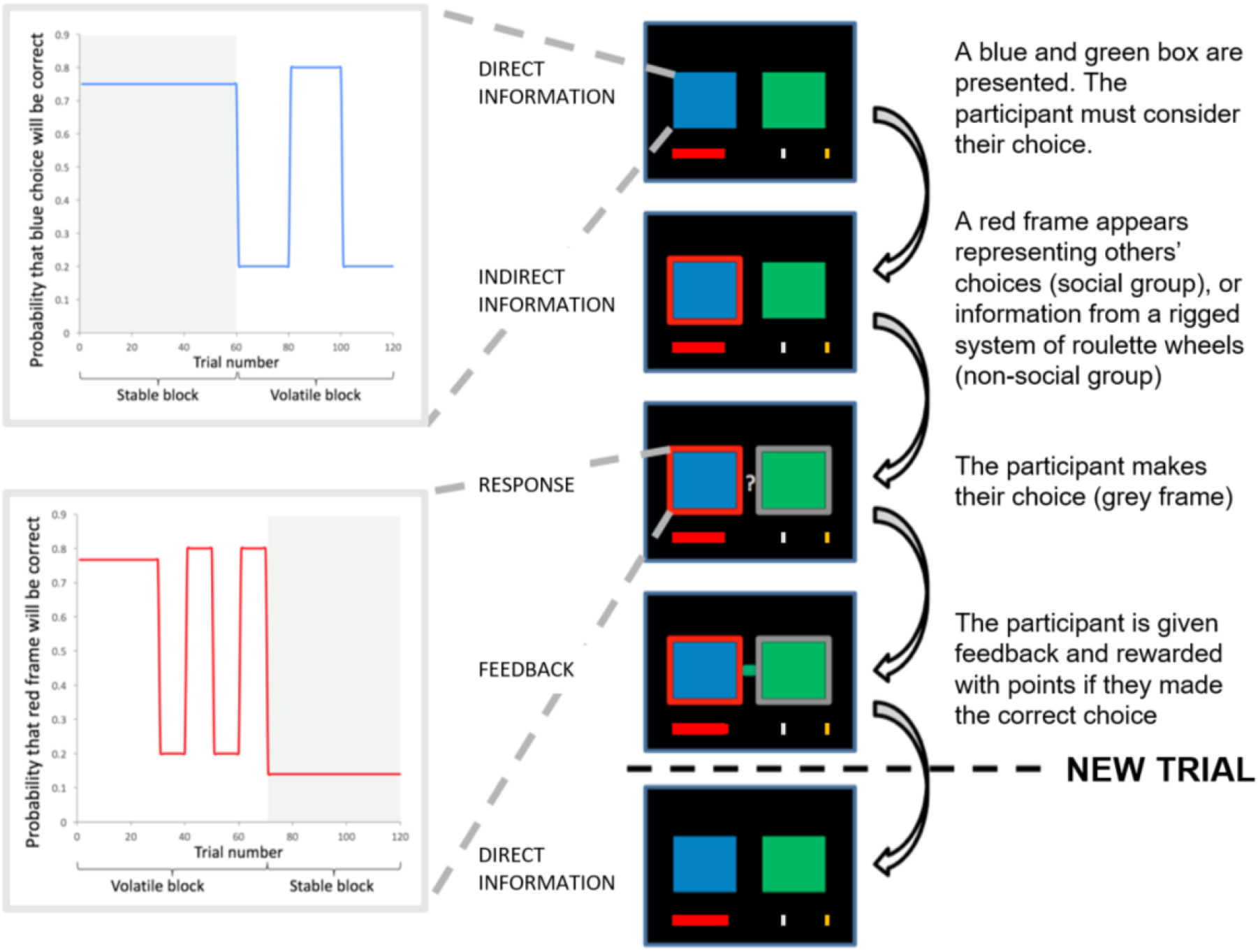
Task Flow Diagram. Participants selected between a blue and green box in order to win points. On each trial, participants saw the direct sources (boxes), then either the blue or the green box was highlighted with a red frame (the indirect source). Participants were instructed that the frame represented either the most popular choice made by a group of participants who had completed the task previously (Social Group) or the choice from rigged roulette wheels (Non-social Group). Immediately after participants had responded, their selected option was framed in grey. A 0.5–2 s interval ensued, after which participants received feedback in the form of a green or blue box in the middle of the screen. If participants were successful, the red reward bar progressed toward the silver and gold goals. The probability of reward associated with the blue and green boxes and the probability that the red frame surrounded the correct box varied according to probabilistic schedules which comprised stable and volatile blocks.

### Social and Non-social Groups exhibit comparable differences in the effect of MPH on direct and indirect learning

#### Lose-shift, win-stay analysis

Win-stay and lose-shift scores were calculated for direct and indirect learning. For indirect learning an example lose-shift trial would be one in which the participant selected the same option as the red frame, lost, and subsequently shifted to selecting the option not surrounded by the red frame. In our task the utility of a win-stay, lose-shift strategy varies according to environmental volatility (e.g. this strategy is adaptive in volatile phases but mal-adaptive in stable phases where optimal performance depends on ignoring misleading probabilistic feedback). We therefore calculated separate scores for volatile and stable phases and included volatility as a factor in our analysis.

To investigate whether effects of MPH varied as a function of the direct versus indirect, or social versus non-social, nature of an information source we submitted win-stay and lose-shift scores to a repeated measures ANOVA with within-subjects factors drug (MPH, PLA), volatility (stable, volatile), directness (direct, indirect) and index (win-stay, lose-shift), and between-subjects factor group (social, non-social). We predicted that, if there is indeed an important distinction between social and non-social learning we should see a significant interaction involving the factors drug and group. If, on the other hand, differential effects of MPH are driven, not by social nature, but by direct versus indirect learning we should see an interaction involving the factors drug and directness, and no interaction with group. Only the former would be considered evidence for separable social-specific mechanisms.

We observed a significant **drug x volatility x directness** interaction (F(1,100) = 7.393, p = 0.008). Simple effects analyses revealed a significant difference between volatile and stable scores under PLA for direct (volatile mean(sterr) 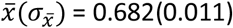, stable 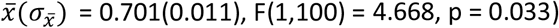, but not for indirect learning 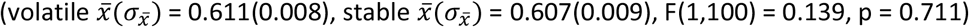, and no difference between volatile and stable scores under MPH for direct 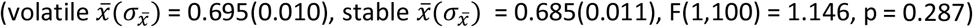, or indirect learning 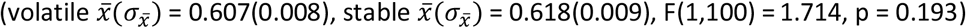. Thus, under MPH participants exhibited more win-stay and lose-shift behaviour in volatile compared to stable environments, whereas the opposite was true under PLA. However, this effect of MPH is restricted to learning from the direct source (the blue and green boxes) and does not extend to learning from the additional indirect information represented by the red frame. To illustrate this interaction, we display in **Figure 2a** the difference between volatile and stable blocks; a high score reflects superior adjustment to block volatility. Note that there was no drug x group (F(1,100) = 0.230, p = 0.633), drug x volatility x group (F(1,100) = 0.039, p = 0.845), drug x volatility x directness x group interaction (F(1,100) = 0.049, p = 0.825) and no other significant interactions involving the factor drug (all p > 0.05). Drug effects as a function of group are illustrated in **Figure 2c**. See **Supplementary Information 3** for main effects and interactions not involving the factor drug and for exploration of drug administration order effects.

**Figure 2.**
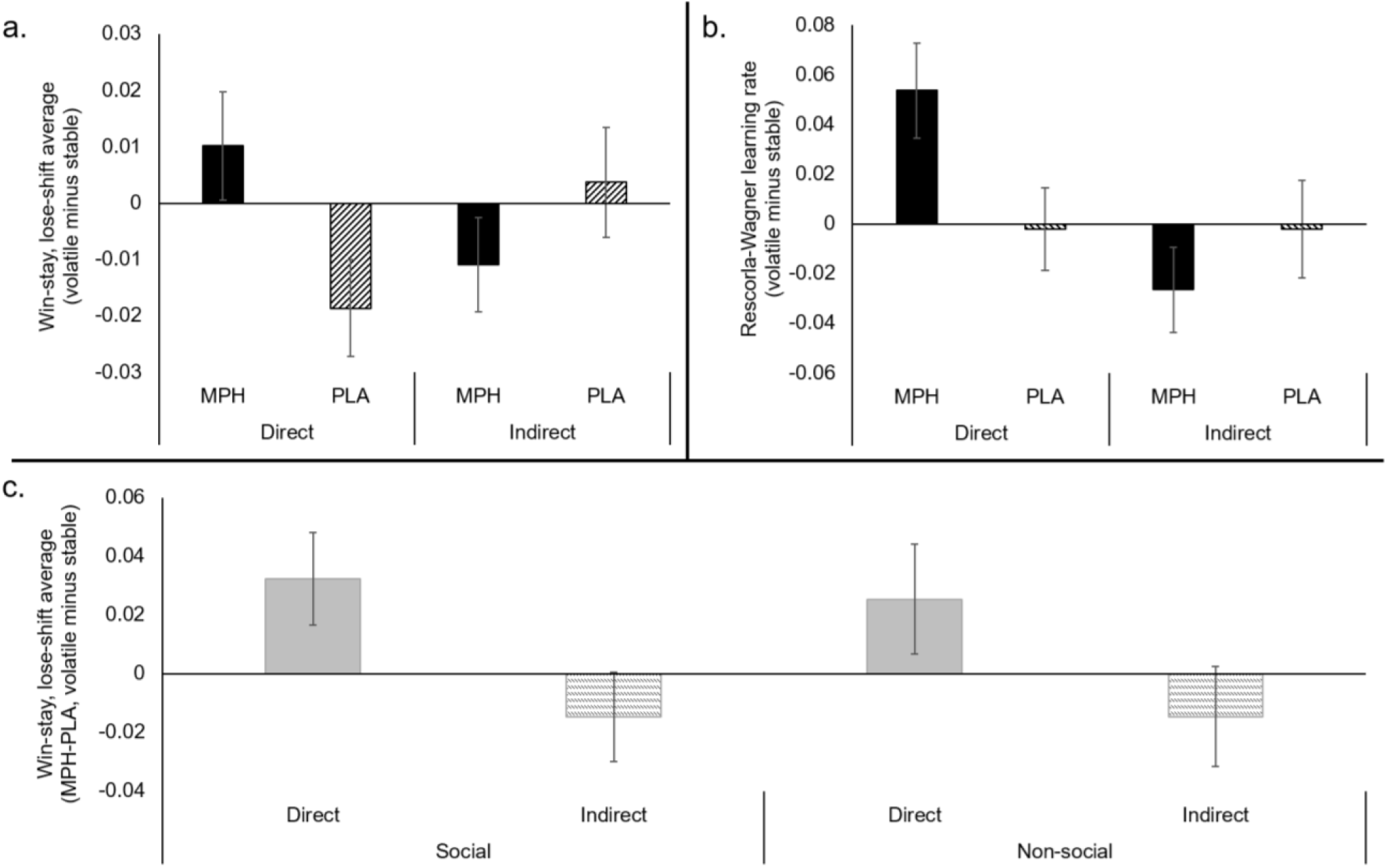
a. Lose-shift and win-stay drug x directness x volatility interaction. Relative to PLA, MPH promoted an adaptive change in lose-shift and win-stay scores for direct, but not for indirect learning (collapsed across group). **b. Learning rate drug x directness x volatility interaction**. Relative to PLA, MPH increased the change in learning rate between volatile and stable blocks for direct, but not for indirect learning (collapsed across group). **c. Comparable drug effects for Social and Nonsocial Groups**. The difference in the effect of MPH on direct and indirect learning was comparable between Social and Non-Social Groups. Error bars represent standard error of the mean.

To provide a final test of whether it is accurate to claim that drug effects did not differ as a function of the social nature of the cover story we computed one single difference score representing the differential effects of MPH on direct and indirect learning in stable and volatile environments. This score was calculated separately for Social and Non-social Groups and submitted to a Bayesian independent samples t-test (https://jasp-stats.org^19^). The BF_10_ value was 0.214 thus providing support for the null hypothesis that the groups do not differ.

#### Rescorla-Wagner analysis

Mathematical models of learning suggest that beliefs are updated in proportion to prediction errors – the difference between predicted and actual outcomes – which are modulated by a learning rate (*α* alpha)^20^. The above win-stay, lose-shift analysis makes the prediction that *α* values relating to direct learning should vary as a function of both drug and volatility. To test this prediction we estimated *α* values by fitting an adapted Rescorla-Wagner learning model^20^ which estimates *α* for direct and indirect learning sources and for volatile and stable learning environments separately and therefore provides four *α* estimates: *α*_stable_direct_, *α*_voLdirect_, *α*_stablejndirect_, *α*_voljndirect_ (see **Supplementary Information 7** for details of model fitting and model comparisons). A repeated measures ANOVA with within-subjects factors drug (MPH, PLA), volatility (stable, volatile), and directness (direct, indirect), and between-subjects factor group (social, non-social) showed a significant **drug x volatility x directness** interaction (F(1,100) = 6.913, p = 0.010) and, crucially, no drug x volatility x directness x group interaction (F(1,100) = 0.435, p = 0.511). Simple effects analyses revealed a significant difference between volatile and stable scores under MPH for direct 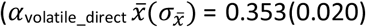, 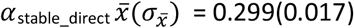, F(1,100) = 8.116, p = 0.005), but not for indirect learning 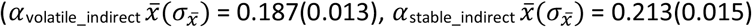, F(1,100) = 2.848, p = 0.095), and no difference between volatile and stable scores under PLA for direct 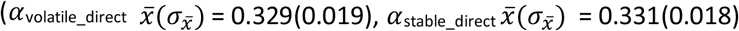, F(1,100) = 0.020, p = 0.888), or indirect learning 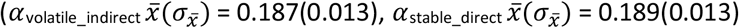, F(1,100) = 0.025, p = 0.876). To illustrate this interaction, we display in **Figure 2b** *α*_vol_direct minus_ *α*_stable_direct_, and *α*_vol_indirect_ minus *α*_stable_indirect_; a high score reflects superior adjustment to block volatility. There were no other significant interactions involving the factor drug (all p > 0.05). See **Supplementary Information 3** for main effects and interactions not involving the factor drug and for exploration of drug administration order effects.

As above, conducting a Bayesian independent samples t-test on scores representing the direct-indirect difference in the MPH-PLA learning rate difference between volatile and stable revealed support for the null hypothesis that the Social and Non-social groups do not differ (BF_10_ = 0.254).

Another influence on choice behaviour, and thus another potential locus of effect of MPH, concerns the process of transforming learned probabilities into action choices. We formally described this process using a softmax action selector model that contains the free parameters beta *(β)* – which controls the extent to which calculations of probability influence choice – and zeta (ζ) – which controls the weighting of indirect relative to direct sources of information. Paired samples t-tests showed no difference in *β* (MPH 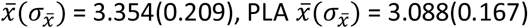; t(101) = 1.152, p = 0.252) or ζ (MPH 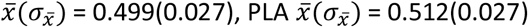; t(101) = −0.554, p = 0.581) values between MPH and PLA conditions.

In sum, MPH increased participants’ ability to adapt to environmental volatility, such that they exhibited increased learning rates under volatile relative to stable conditions but this was only the case for direct learning – from one’s own personal experience. MPH did not modulate participants’ ability to adapt to the volatility of the utility of an indirect source of information.

### MPH selectively affects direct learning in keen indirect learners

The red frame is a highly salient and behaviourally relevant cue; optimal performance on this task requires participants to learn about the utility of the frame and use this information to help choose amongst the boxes. Thus, the absence of an effect of MPH on indirect learning is particularly striking. To guard against the possibility that the lack of effect of MPH on indirect learning is driven by reduced indirect, relative to direct, learning (i.e. participants learn primarily from the blue/green boxes and ignore the frame) we ran a post-hoc analysis. First, we identified whether direct or indirect learning was the primary driver of responses for each of our participants. This was achieved by regressing a Bayesian Learner Model^21^ of optimal responses against each participant’s behaviour under PLA using the same method employed by Cook et al.^10^. This produced two beta vales for each participant representing the extent to which their responses were driven by direct (*β*_direct_) and indirect learning (Indirect). We then selected participants for whom *β*_indirect_ > *β*_direct_ (N = 46). Under PLA, these participants primarily use the indirect information (the frame) to make their decisions. A repeated measures ANOVA with within-subjects factors drug (MPH, PLA), volatility (stable, volatile) and directness (direct, indirect) showed a significant drug x volatility x directness interaction (F(1,45) = 13.338, p = 0.001). Under MPH *α*_vol_direct_ was significantly greater than *α*_stable_direct_ 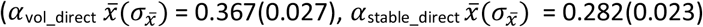, F(1,45) = 12.081, p = 0.001) but not under 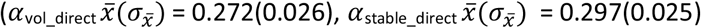, F(1,44) = 0.587, p = 0.448). For indirect learning there was no interaction between drug and volatility and no main effect of drug (all p > 0.05). Thus, we observed a selective effect of MPH on direct learning even for participants whose responses under PLA are primarily driven by indirect learning.

### No group differences for participants particularly sensitive to the effect of MPH on indirect learning

To guard against the possibility that the lack of a group difference is influenced by an insensitivity of indirect learning to the effects of MPH, we selected a subgroup of participants who were particularly sensitive to the effects of MPH on indirect learning ((MPH-PLA *α*_stablejndirect_ + MPH-PLA *α*_volatile_indirect_)>(MPH-PLA *α*_stable_direct+_ MPH-PLA *α*_volatile_direct_)). For this subgroup (Social Group N = 26, Non-social Group N = 27) we ran a repeated measures ANOVA with within-subjects factors drug (MPH, PLA), volatility (stable, volatile), and directness (direct, indirect) and between-subjects factor group (social, non-social). The interaction between drug, volatility and directness remained significant (F(1,51) = 5.506, p = 0.023). There was no drug x volatility x directness x group interaction (F(1,51) = 0.813, p = 0.371), no main effect of group (F(1,51) = 1.415, p = 0.240) and no other significant interactions involving group (all p > 0.05; Bayesian t-test comparing groups on scores representing the direct-indirect difference in the MPH-PLA *α* difference between volatile and stable BFio = 0.386). As above, under MPH 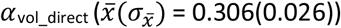 was greater than 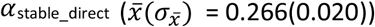, though this difference only approached significance F(1,51) = 3.474, p = 0.068). Under PLA 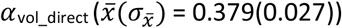 and 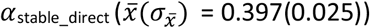 were comparable (F(1,51) = 0.714, p = 0.402). There were no differences between *α*_vol_indirect_ and *β*_stable_indirect_ under MPH (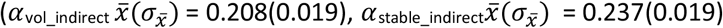, F(1,51) = 2.559, p = 0.116), or PLA (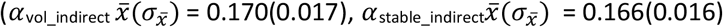, F(1,51) = 0. 060, p = 0.807).

### MPH selectively affects direct learning in the Social Group

As highlighted in the above introduction, the view that there is something special about *social* learning is highly persistent and has been supported by studies demonstrating that social and individual learning are at least partially dissociable. Indeed using a similar task to that employed here, Behrens and colleagues demonstrated activity relating to social and individual learning in dissociable neural pathways^8^. Consistent with this, if we <u>focus exclusively on the Social Group</u>, it appears that MPH improves individual learning (i.e. direct learning) from the boxes but not social (indirect) learning from the red frame (drug x volatility x directness interaction: Win-stay, lose-shift analysis F(1,49) = 5.064, p = 0.029, Rescorla-Wagner analysis F(1,49) = 4.850, p = 0.032). However, as demonstrated in our main analysis, this pattern of data is also observed in the Non-social Group, and Bayesian statistics provide support for the null hypothesis that the Social and Non-social groups do not differ. Thus, it is unlikely that the lack of effect of MPH on learning from the frame is related to the social nature of the information that the frame represents, and more likely that it is related to the demand for indirect learning.

### Summary

In sum, relative to PLA, MPH modulated learning from the direct source of information (the blue and green boxes) as indexed by increased a) win-stay and lose-shift behaviour, and b) learning rate, in volatile compared with stable conditions. If one focuses exclusively on the Social Group, MPH improves individual learning from the boxes but not social learning from the frame and therefore dissociates the two learning types. However, by including a group who received a non-social cover story we have demonstrated that selective effects of MPH are better explained by direct versus indirect, rather than by social versus non-social, learning.

## DISCUSSION

In line with the view that humans possess adaptive specialisations for social learning, previous studies have suggested that social and individual learning are at least partially dissociable. However, in most extant studies, the social source is also an additional, indirect, source of information, thus its nature (social versus non-social) and directness (indirect versus direct) are confounded. Here we assessed whether any dissociable effects of a catecholaminergic challenge on individual versus social learning are best explained by social versus non-social, or direct versus indirect, learning.

Participants completed a complex decision-making task, requiring them to concurrently learn from their own personal experience, and from an additional indirect source of information represented by a red frame. One group was told that the frame represented social information, a second group received a non-social cover story about the frame. Participants completed the task once under PLA and once after MPH. Strikingly, we found that although the effect of MPH varied as a function of direct versus indirect learning, there were no differences between Social and Non-social groups. In other words, with respect to direct learning (feedback-based learning about the blue and green boxes) MPH improved the ability to adapt to changes in the learning environment. However, MPH had no effect on learning from the additional indirect source of information (frame) and this was true irrespective of whether the indirect source was believed to be social or non-social in nature.

Our study cannot provide evidence in support of the adaptive specialisation view: that is, our study did not shown a difference in the effect of drug as a function of group (social / non-social). Instead, our pattern of data suggests that dissociable effects of MPH are driven by direct versus indirect, and not by social versus non-social, learning.

Although our results provide evidence for a neurochemical dissociation between learning from direct and indirect sources of information, we find no evidence for a dissociation between social and non-social learning. Note that the lack of a difference between social and non-social groups cannot be due to poor comprehension of the cover story: participants were prohibited from starting the task until they scored 100% correct in a quiz which included questions about the nature of the red frame (see **Methods: learning task**). Furthermore, using the same task, we have previously demonstrated that social dominance predicts learning for participants that are given the social, but not the non-social, cover story^10^. Thus, belief about the origin of the information represented by the frame modulates the way in which this information is processed. How should our results be interpreted in the context of previous work arguing for adaptive specialisations for specifically social learning^6,7^? It is clear that our results do not provide evidence in support of the adaptive specialisation view. Evidence in support of this view would have come in the form of a social/nonsocial group difference (e.g. enhanced learning under MPH for all learning types except for social learning (i.e. direct learning and indirect learning from a non-social source)). Our Bayesian analysis supports the conclusion that the effects of MPH on learning from social and non-social information are comparable. One could argue that, in addition to failing to support the adaptive specialisation view, our results conflict with this view because they raise the possibility that social and non-social learning share overlapping (neurochemical) mechanisms. This suggestion is consistent with a number of recent studies which have largely failed to find convincing neural evidence for social-specific learning mechanisms. These studies have instead suggested that social learning, like nonsocial learning, is governed by the domain-general process of comparing predictions with incoming sensory evidence, computing prediction errors and using these prediction errors to refine future predictions^1^. Indeed, blood oxygenation level dependent (BOLD) signals correlating with these ‘social prediction errors’ have been reported in brain areas classically associated with non-social learning including the striatum and orbitofrontal cortex^2,21–25^ Nevertheless, it is possible that social and non-social learning are underpinned by networks that can be neurochemically dissociated, though by a pharmacological agent other than MPH (e.g. by a serotonergic agent^e.g.27,28^); this possibility should be explored in future studies.

The absence of an effect of MPH on indirect learning is particularly striking given that the red frame is a highly salient, behaviourally relevant, attentional cue. Why would MPH selectively affect direct, but not indirect learning? One possibility is that our participants simply ignored the information that was represented by the frame; in other words, the current paradigm is poorly suited to detecting variation in indirect learning. In opposition to this, we have demonstrated that MPH selectively affects direct learning *even for participants whose responses are primarily driven by indirect learning* (see results section ‘MPH selectively affects direct learning in keen indirect learners’). Thus, the lack of effect of MPH on indirect learning is not driven by a tendency to ignore the frame.

Another possible explanation for direct learning-selective effects of MPH relates to the complex effects of MPH on working memory: Fallon and colleagues^29^ have demonstrated that whereas MPH improves distractor resistance it impairs working memory updating. More specifically, in a delay period between initial presentation of to-be-remembered information and recall, Fallon and colleagues presented additional information which participants should sometimes ignore and sometimes use to replace (update) the original to-be-remembered information. MPH improved ignoring, but not updating – an effect which the authors interpret in the context of dual state theory^30^ which postulates that intermediate levels of dopamine favour stabilisation of representations whereas excessively high or low levels of dopamine favour representation destabilisation. The current task can be conceptualised in a similar fashion to Fallon and colleagues’ paradigm: participants first see the direct source (boxes) for a variable delay which provides the opportunity to reflect on the history of outcomes and estimate the value associated with each box. In the ‘delay period’ between seeing the boxes and making their response, participants see the indirect source (frame). By reflecting on the history of the veracity of the frame, they can estimate the probability that the frame provides correct advice, and decide whether to <u>update</u> their choice, or <u>ignore</u> the frame. Positive effects of MPH on ignoring, not updating, would result in a benefit to direct learning – i.e. direct learning should be more robust against misleading information from the frame – but not indirect learning because improved distractor resistance benefits the first information source (boxes) but does not benefit the information source that is presented in the delay period. Thus, the complex effects of MPH on working memory provide a potential explanation for the direct learning-selective effects of MPH observed here.

Our results not only show a direct-learning selective effect of MPH, but also demonstrate a selective effect on (direct) learning about the volatility of the environment. This finding that MPH, which blocks both noradrenaline and dopamine transporters, improves adaptation of learning rates to reflect environmental volatility, fits well with existing literature which documents a role for noradrenaline in flexibly adapting to environmental change. Theoretical and modelling work has suggested that noradrenaline acts as a ‘neural interrupt signal’ which alerts the learner to an unpredicted change in the learning environment^31,32^. In line with this, pharmacological manipulations and lesions of the noradrenergic system affect performance, in non-human animals, in tasks such as reversal learning and attentional-set shifting which require detection of and adaptation to environmental changes^33–39^. In humans, learning rates^40–42^ and prediction errors^43,44^ are correlated with pupil size, an indirect index of locus coeruleus (the main noradrenergic nucleus of the brain) activity^45–48^, and are modulated by pharmacological manipulation of the noradrenaline system^15,16^. In the context of our task, the ‘unpredicted change in learning environment’ corresponds to a switch from stable to volatile environment. By administering MPH to our participants we have likely increased their synaptic noradrenaline levels^49^ and reduced spontaneous activity in locus coeruleus neurons via noradrenaline action at α-adrenergic receptors on locus coeruleus cells^50^. Thus, it is likely that MPH has shifted locus coeruleus activity toward a phasic mode, in which cells respond more strongly (and specifically) to salient events^51^ such as the switch from stable to volatile environments. Consistent with the neural interrupt hypothesis we found that learning rates were better adapted to suit environmental volatility under MPH relative to PLA.

Dopamine has also been implicated in the regulation of learning rate^52^. Individual differences in human learning rates have been associated with genetic polymorphisms coding for the catechol-O-methyltransferase enzyme which plays a role in metabolizing dopamine, the dopamine transporter, and D2 receptors^53,54^. A role for dopamine in regulation of learning rate is also consistent with studies indicating that dopamine neurons respond to novel and unexpected stimuli, and that dopamine is important for cognitive flexibility^55,56^. It should, however, be noted that studies that have tried to disentangle the roles of dopamine and noradrenaline in complex learning tasks have argued that, while noradrenaline is associated with ‘higher level’ learning about environmental change, dopamine is linked to the ‘lower level’ modulation of motor responses as a function of learned probabilities^16^. Given this, and the known effects of MPH on synaptic dopamine^17^, it is somewhat surprising that we observe selective effects of MPH on higher level learning about environmental volatility, in the absence of a main effect of drug on learning rates. This pattern of results is reminiscent of the effects of MPH on intra-dimensional/extra-dimensional set shift tasks^57^. In such tasks participants learn a discrimination on the basis of a particular stimulus dimension (e.g. colour: blue = reward, green = no-reward); participants are subsequently subjected to intra-dimensional shifts, wherein the same dimension is relevant but reward contingences are reversed (blue = no-reward, green = reward), and extra-dimensional shifts wherein a stimulus dimension (e.g. shape) that was initially irrelevant is now relevant. Thus, the task indexes (a) ‘higher level’ learning – attending to environmental cues relevant for reinforcement (extra-dimensional shift); (b) ‘lower level’ learning – relearning of previously acquired stimulus-reward associations (intra-dimensional shift). Consistent with our results, Rogers and colleagues^57^ demonstrated that MPH improved higher level (extra-dimensional), but not lower level (intra-dimensional) learning. One might hypothesise that these effects of MPH on adaptation to environmental volatility are primarily mediated by noradrenergic, rather than dopaminergic, function. However, further studies which might, for example, administer MPH in conjunction with selective dopamine and/or noradrenaline receptor blockers, are needed to test this hypothesis.

The absence of an effect of MPH on indirect learning from an additional source even when the indirect source is believed to be social in nature is perhaps surprising given positive effects of MPH on social influence^58^, reports that social learning activates catecholamine rich areas of the brain such as the striatum^2,23,25,26,59,60^, and evidence that ventral striatal prediction errors during social learning are influenced by a gene involved in regulating dopamine degradation^2^. An important consideration, however, is that in studies that have linked social learning to signals in catecholamine-rich areas of the brain, participants have not been required to learn from their own personal experience. Thus, the social information is not an *additional* source to be learned about in concert with learning from one’s own experience, rather, the social source is the *only* source that participants must learn about. Contrasting our study with this existing literature makes the concrete prediction that, if roles could be reversed such that social information comprises the direct source and individual information comprises the indirect source, one would observe a significant effect of MPH on social, but not individual, learning.

## Conclusion

In most extant paradigms that have dissociated social and individual learning, social information comprises an indirect, additional, source and thus differs from individual learning both in terms of social nature (social versus non-social) and directness (indirect versus direct). Here we provide evidence that selective effects of MPH are better explained by direct versus indirect, rather than by social versus non-social, learning. Specifically, we showed that relative to PLA, MPH modulated learning from a direct source of information (blue and green boxes) as indexed by increased a) win-stay and lose-shift behaviour, and b) learning rate, in volatile compared with stable conditions. MPH did not modulate learning from an indirect source (frame), and this pattern of data was comparable for Social and Non-social Groups. These data fail to support the adaptive specialisation view which postulates separate mechanisms for social and non-social learning, instead providing positive evidence for distinct mechanisms for direct versus indirect learning.

## METHODS

The general method is identical to that reported in Swart et al^61^ and is included here for completeness.

### General procedure and pharmacological manipulation

The study consisted of two test sessions with an interval of one week to 2 months. The first test day started with informed consent, followed by medical screening. Participation was discontinued if participants met any of the exclusion criteria. On both test days participants first completed baseline measures. Next participants received a capsule containing either 20 mg MPH (Ritalin®, Novartis) or PLA, in a double-blind, placebo-controlled, cross-over design. MPH blocks the dopamine and noradrenaline transporters, thereby diminishing the reuptake of catecholamines. When administered orally, MPH has a maximal plasma concentration after 2 hours and a plasma half-life of 2-3 hours^62^. After an interval of 50 minutes, participants started with the task battery containing the learning task. On average the learning task was performed 1 hour 20 minutes after capsule intake, well within the peak of plasma concentration. Both test days lasted approximately 4.5 hours, which participants started at the same time (maximum difference of 45 minutes). Blood pressure, mood and potential medical symptoms were monitored thrice each day: before capsule intake, upon start of the task battery and after finishing the task battery. Participants were told to abstain from alcohol and recreational drugs 24 hours prior to testing and from smoking and drinking coffee on the days of testing. Participants completed self-report questionnaires at home between (but not on) test days. Upon completion of the study, participants received monetary reimbursement or study credits for participation. The study was in line with the local ethical guidelines approved by the local ethics committee (CMO protocol NL47166.091.13; trial register NTR4653) and in accordance with the Helsinki Declaration of 1975. Baseline measures, self-report questionnaires, mood-ratings and medical symptoms are reported in **Supplementary Information 4**.

### Participants

Data was collected from 106 native Dutch volunteers (aged 18 – 28 years, 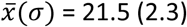; 53 women; 84 right-handed). Four participants dropped out after the first test day (due to too much delay between test days, loss of motivation, nausea, and mild arrhythmia). Of the resulting 102 participants, 50 participants received MPH on the first day. Exclusion criteria comprised a history of psychiatric, neurological or endocrine disorders. Further exclusion criteria were autonomic failure, hepatic, cardiac, obstructive respiratory, renal, cerebrovascular, metabolic, ocular or pulmonary disease, epilepsy, substance abuse, suicidality, hyper/hypotension, diabetes, pregnancy/breastfeeding, lactose intolerance, regular use of corticosteroids, use of psychotropic medication or recreational drugs 24 hours before each test day, and first degree family members with schizophrenia, bipolar disorder, ventricular arrhythmia or sudden death. Excluded volunteers received €10,-as a compensation for their time and effort.

### Learning task

Participants completed a modified version^10^ of a decision making task developed by Behrens and colleagues^8^. On each trial participants made a choice between a blue and a green box. Correct choices were rewarded with points represented on a bar spanning the bottom of the screen. Participants’ aim was to accumulate points to obtain a silver (€2) or gold (€4) reward. Throughout the experiment the probability of reward associated with each option (blue or green) varied according to a probabilistic schedule. Participants were informed that the task followed ‘phases’ wherein sometimes blue, but at other times green, would be more likely to result in reward.

In addition to the reward history, a second source of information was available to participants. On each trial, before participants made their choice, a frame appeared. Participants in the Social Group were instructed that this frame represented choices made by a group of participants who had completed the task previously. Participants in the Non-social Group were instructed that the frame represented the outcome from a system of rigged roulette wheels. Both groups were informed that the frame would fluctuate between providing predominantly correct and predominantly uninformative ‘advice’ (see **Supplementary Information 2** for instruction scripts).

Participants in both the Social and Non-social Groups were pseudorandomly allocated to one of four different randomisation groups, group membership determined the probabilistic schedule underpinning outcomes (blue/green) and the veracity of the frame (correct/incorrect) (see Supplementary Information 1 for further details and for statistical analyses including randomisation group as a factor). Although probabilistic schedules for Day 2 were the same as Day 1 there was variation in the trial-by-trial outcomes and advice. To prevent participants from transferring learned stimulus-reward associations from Day 1 to Day 2 different stimuli were also employed: 50% of participants viewed yellow/red triangles with advice represented as a purple frame on Day 1 and blue/green squares with advice as a red frame on Day 2, for 50% of participants this order was switched. Before participation all participants completed a self-paced step-by-step on-screen explanation of the task in which they first learned to choose between two options to obtain a reward, and subsequently learned that the advice represented by the frame could help them in making an informed decision about their choice. Following the task explanation, participants were required to complete a short quiz testing their knowledge of the task. Participants were required to repeat the task explanation until they achieved 100% correct score in this quiz. Consequently, we could be sure that all participants understood the structure of the task and were aware that the red/purple frame represented either social information (Social Group), or information from a rigged set of roulette wheels (Non-social Group).

All participants completed 120 trials on Day 1 and Day 2. The task lasted approximately 25 minutes, including instructions. At the end of the task participants were informed whether they had scored enough points to obtain a silver / gold award.

### Statistical analyses

*Win-stay, lose-shift analysis*

For direct learning a trial was denoted as a win-stay if the participant had won on the previous trial and if their response on the current trial was the same (e.g. they selected blue and won on the previous trial, and subsequently re-selected blue). A trial was denoted as a lose-shift if the participant had lost on the previous trial and if they subsequently shifted their response. Win-stay and lose-shift trials were summed and divided by the total number of win/lose trials to control for differing numbers of win/lose trials between randomisation schedules and (volatile/stable) blocks. Shapiro-Wilk tests showed that averaged win-stay and lose-shift scores, for direct and indirect learning, volatile and stable conditions, and for social and non-social groups, did not significantly deviate from the normal distribution (see **Supplementary Information 5** for table of statistics). Win-stay and lose-shift scores were entered into a single repeated measures ANOVA with within-subjects factor drug (MPH, PLA), volatility (stable, volatile), directness (direct, indirect) and index (win-stay, lose-shift).

### Rescorla-Wagner analysis

Learning rates for stable and volatile blocks were calculated by fitting a simple learning model to participants’ choice data. This learning model consisted of two Rescorla-Wagner predictors^20^ (one which estimated the utility of a blue choice (*r__direct(i+1)_*) and one which estimated the utility of going with the red frame (*r__indirect(i+1)_*)) coupled to an action selector^21,40^. This was achieved using the scripts tapas_rw_social_reward_vol.m and rw_softmax_constant_weight_social_reward.m which are available at https://github.com/JenniferLcook/sociallearning, and tapas_quasinewton_optim.m and a modified version of tapas_fitModel.m which are freely available as part of the open source software package TAPAS at http://www.translationalneuromodeling.org/tapas. See **Supplementary Information 6** for details of the model, model fitting, and for evaluation of the model’s performance (ability to account for participant choice), see **Supplementary Information 7** for model comparisons.

Rescorla-Wagner predictors:

Our Rescorla-Wagner predictors comprised modified versions of a simple delta-learning rule^20^. This rule typically has a single free parameter, the learning rate *(α)* and estimates outcome probabilities using equation (1).

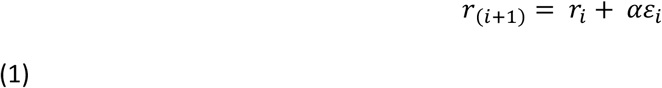

where *r_(i+1)_* is the predicted outcome probability for the (*i*+1)th trial, *r_i_* is the predicted outcome probability for the ith trial, *ε_i_* is the prediction error at the ith trial, and *α* is the learning rate. By choosing different values for *α*, the model can make different approximations of a participant’s outcome probability estimates. Our modified version separately estimated *α* for stable and volatile blocks. Furthermore, we simultaneously ran two Rescorla-Wagner predictors such that we could estimate *α* values relating to both direct and indirect learning. Subsequently, our model generated the predicted outcome probability of a blue choice inferred from previous experience (*r__direct(i+1)_*) and of the choice to go with the red frame (*r__ndirectβi+1_*)) and provided four *α* estimates: *α_stable_direct_, β_vol_direct_, α_stable_indirect_, β_vol_indirect_*.

Action selector:

As participants had access to both information from direct learning and indirect learning our response model assumed that participants integrated both sources of information in order to predict the probability that a blue choice would be rewarded. Based on work by Diaconescu and colleagues ^1,2^ we calculated the estimated value of a blue choice according to equation (2):

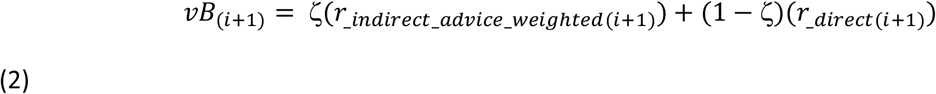

In equation (2), ζ is a parameter that varies between individuals and which controls the weighting of indirect relative to direct sources of information. *r__indirect advice weighted(i+1)_* comprises the advice provided by the frame weighted by the probability of advice accuracy (*r__ndirect(o+1)_*) in the frame-of-reference of making a blue choice (equation (3)).

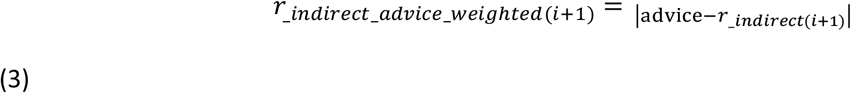

where advice = 0 for blue and advice = 1 for green. Thus, if the frame advised ‘green’ and the probability of advice accuracy was estimated at 80% (*r__indirect(i+1)_* = 0.80), the probability, inferred from indirect learning, of a blue choice being rewarded would be 0.2 (*r__indirect_advice_weighted(i+1)_* = 11 – 0.8| = 0.2). Conversely the probability, inferred from indirect learning, that the correct choice is green would equal 0.8.

The probability that the participant follows this integrated belief (*υB_(i+1)_*), was described by a sigmoid function (equation (4)); here, responses are coded as *y*_(*i*+1)_ = 1 when selecting the blue option, and *y_(i+1)_* = 0 when selecting green.

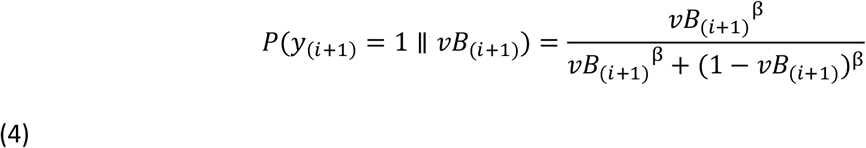

β is a participant-specific free parameter representing the inverse of the decision temperature: as *υB*_(*i+1)*_, the sigmoid function approaches a step function with a unit step at *υB*_(*i*+1)_= 0.5 (i.e., no decision noise).

### Bayesian statistical analysis

Results were also analysed within a Bayesian framework using JASP^19^ to examine the strength of the evidence in favour of the null and experimental hypotheses with a prior of a medium effect size using default priors. Bayes Factors provide a ratio of the likelihood of the observed data under the null versus alternative hypothesis, whereas *p*-values examine the probability of the data given the null hypothesis and therefore cannot discriminate between evidence for the null and no evidence for either the null or alternative hypothesis^63^. Bayes Factors (BF_10_) are reported. By convention, values < 1/3 and > 3 are taken as evidence in favour of the null and alternative hypotheses, respectively, while values within these boundaries are judged to provide no evidence to favour either the null or alternative hypotheses. We computed one average lose-shift, win-stay difference score representing the differential effects of MPH on direct and indirect learning in stable and volatile environments (e.g. the direct-indirect difference in the MPH-PLA difference between volatile and stable), this score was calculated separately for Social and Non-social Groups and submitted to a Bayesian independent samples t-test; BF_10_ > 3 would therefore provide support for the alternative hypothesis that the Social and Non-social Groups differ, BF_10_ < 0.33 would provide support for the null hypothesis that the Social and Non-social Groups are comparable. The same analysis was also run on difference scores representing the differential effects of MPH on direct and indirect learning in stable and volatile environments for Rescorla-Wagner *α* values.

**Table 1.** There were no significant differences between the social and non-social groups in age, gender or socioeconomic status as measured using the Barratt Simplified Measure of Social Status (Barratt, 2012).

|  | Social | Non-social |  |
| --- | --- | --- | --- |
| Age ($\bar{x}(\sigma)$) | 21.780(2.582) | 21.289(2.022) | $t(100) = 1.068, p = 0.288$ |
| Male:Female | 26:24 | 25:27 | $\chi^2(1) = 0.157, p = 0.692$ |
| Socioeconomic status ($\bar{x}(\sigma)$) | 60.359(7.771) | 59.219(8.378) | $t(100) = 0.504, p = 0.615$ |
| N | 50 | 52 |  |

## Acknowledgements

We thank Dr Monique Timmer and Dr Peter Mulders for medical assistance, and Dr Sean James Fallon for advice on setting up the MPH study.

## Author contributions

JC, RC, HdO designed the study. JC, HdO, JS, MJF and DG collected the data. JC, AD and JS analysed the data. JC, JS, and RC drafted the manuscript. All authors edited the manuscript.

